# Signal, noise, and sampling: How pool size and replication shape metabolomic inference

**DOI:** 10.64898/2026.04.07.717001

**Authors:** David L. Hubert, Dylan L. Porter, Ryan D. Robinson, Maya E. Mijares, Elmira Ahmadian, Kenneth R. Arnold, Mark A. Phillips

## Abstract

Metabolomics provides a direct readout of physiological state and is increasingly used in evolutionary and systems biology. In small organisms such as Drosophila melanogaster, metabolomic analyses typically require pooling individuals to obtain sufficient material, yet pool sizes vary widely across studies with little justification. How pooling and biological replication influence metabolome characterization and the detection of biological signal remains poorly understood.

Here, we evaluate the effects of pool size and biological replication on metabolomic profiles and signal detection using two complementary experimental designs. In the first, we assess how pooling (5, 50, or 100 individuals) influences metabolomic structure and reproducibility in inbred and outbred populations. In the second, we test how pool size interacts with systematic variation in replicate number to affect detection of diet-associated metabolite changes under a high-sugar perturbation.

Pool size strongly influenced metabolomic profiles, with samples pooled at five individuals consistently differing from larger pools, while profiles from 50 and 100 individuals were more similar. Larger pools improved reproducibility in a dataset-dependent manner. In the dietary experiment, smaller pool sizes substantially reduced sensitivity, leading to loss of true diet-associated metabolites without increasing false discoveries. Replicate downsampling further revealed that both pool size and biological replication jointly determine signal retention, with smaller pools accelerating the loss of detectable metabolites under reduced replication.

Across all analyses, the ability to detect metabolite signals was strongly dependent on effect size and variability. Metabolites with larger and more stable effect estimates were consistently retained, whereas those with smaller or more variable effects were rapidly lost under reduced sampling. Linear mixed-effects modeling confirmed that detection probability is governed by a balance between biological signal strength and measurement variability, with pool size and replication jointly modulating this relationship.

More broadly, our results demonstrate that metabolomic inference is governed by the interplay of signal, noise, and sampling design, with pool size and replication jointly shaping the detectability, stability, and interpretation of biological signals.

## Introduction

Metabolomics, the comprehensive analysis of small-molecule metabolites in biological systems, offers a functional readout of cellular physiology (Johnson et al., 2016). The metabolome serves as a bridge between genotype and phenotype, reflecting the combined influence of genetic variation, environmental conditions, and organismal state (Nicholson et al., 1999). Because metabolites act as building blocks, enzyme cofactors, and mediators of energy production and signaling, metabolomic profiling provides a direct window into the molecular processes underlying biological function. Advances in high-resolution mass spectrometry and computational analysis have made it possible to characterize metabolite profiles at increasingly large scales, expanding the use of metabolomics across physiology, disease biology, and evolutionary studies (Clish, 2015; Tautenhahn et al., 2012).

*Drosophila melanogaster* has become a central model for this work due to its short generation time, tractable genetics, and long history in experimental biology. Integrating metabolomics with genetic and systems-level approaches has revealed substantial variation in metabolic architecture across lines and populations, demonstrating how genetic background shapes physiological traits (Ayroles et al., 2009; Mackay et al., 2012; Zhou et al., 2020). Metabolomic profiling has also been incorporated into experimental evolution studies, where replicated populations evolving under defined conditions allow direct links to be drawn between selection, environment, and physiological change (Phillips et al., 2022; Erkosar et al., 2023; Hubert et al., 2025). However, despite the increasingly common use of metabolomics in Drosophila studies, a basic aspect of experimental design remains highly variable across studies: the number of individuals pooled per sample.

As individual flies yield limited material, pooled sampling is standard practice. However, pool sizes differ widely in the literature, ranging from only a few individuals to several dozen or more (e.g., Zhao et al., 2022; Hubert et al., 2025; Vecchié et al., 2026), often without clear justification. This variation introduces an underappreciated source of heterogeneity. Smaller pools may be more sensitive to stochastic differences among individuals or may fail to capture low-abundance metabolites, whereas larger pools may provide more stable estimates but at increased cost and reduced throughput. As metabolomics becomes more central to comparative and evolutionary work, understanding how pooling decisions influence both metabolome characterization and the detection of biological signal is increasingly important.

Here, we evaluate how pooled sample size affects metabolomic characterization and signal detection across two complementary experimental contexts in *D. melanogaster*. In the first, we tested the effect of pool size (5, 50, or 100 individuals) in both inbred and outbred populations sampled at two adult ages. Here outbred populations capture substantial standing genetic variation, whereas inbred populations provide a reduced-variance baseline, allowing us to evaluate how genetic heterogeneity interacts with sampling scale. This design allows us to assess how genetic background influences the degree to which larger pools improve the stability and consistency of metabolomic measurements.

In a second experiment, we focused on inbred populations to test how pool size interacts with replication and treatment effects under a defined dietary perturbation. Replicate populations were maintained on either control or high-sugar diets, and metabolomic profiles were again generated across a gradient of pool sizes within each replicate population. We then systematically downsampled the number of replicates and evaluated metabolomic signal retention across pool sizes at each iteration. This design allows us to evaluate how both pooling and biological replication influence the ability to detect consistent metabolomic differences associated with diet treatment, and to quantify the trade-offs between increasing pool size versus increasing the number of replicates. Together, these experiments address a practical but largely unresolved question in metabolomics: how sampling decisions shape the stability of metabolomic measurements and the power to detect biologically meaningful signals.

## Methods

### Study System

For this experiment, two distinct *Drosophila melanogaster* populations representing different genetic backgrounds were used: inbred and outbred flies. In Experiment 1, the outbred populations were maintained on standard food and used as a laboratory outbred population for approximately 35 generations at the time of this work (CRB). These populations were created by combining 10 females and 5 males form 100 lines from the Drosophila Genetic Reference panel (Mackay et al., 2012; Supplementary Table 1). These lines were derived from a signal natural population in Raleigh, North Carolina. Combining lines effectively creates outbred populations that, at least in part, capture the variation of this source natural population. The inbred population was a OR wild-type (ORWT) strain obtained from Carolina Biological Supply Company (Burlington, NC, USA). In Experiment 1, both populations were maintained on a 28 day life cycle, where eggs were collected, transferred to vials, and incubated for 14 days, then freshly enclosed adult flies were placed into cages for 14 days before the next egg collection was conducted. All populations were provided with identical standard food replaced every 2 days, and were kept under standard laboratory conditions. To minimize environmental variation, cage positions on the shelf were rotated whenever food was replaced to account for any differences in light or temperature.

In Experiment 2, to test how pool size interacts with biological replication and diet treatment to influence metabolomic profiles in an inbred background, we performed a dietary perturbation experiment using the ORWT inbred Drosophila melanogaster strain. Replicate ORWT populations were founded from the laboratory stock and maintained under two dietary conditions: standard food containing approximately 7% molasses (STD) and high-sugar food containing 20% molasses (HSD). Eight independent replicate populations were maintained per diet treatment. Each replicate population comprised 40 vials containing 50–80 adult flies per vial. Flies were collected on their assigned treatment and kept in the same vials in the incubator until sampling to minimize microenvironmental variation. Incubator conditions were 23 °C, 12 h light: 12 h dark.

### Sample Collection

For Experiment 1, technical replicates were generated from a single population from each genetic background (CRB and ORWT) and expanded into three independent replicate populations each. These replicate populations were allowed to develop and reproduce, and eggs collected from each were used to establish experimental cohorts. Samples for metabolomic analysis were collected at two time points, 14 and 40 days post-egg deposition, representing distinct young and old adult stages associated with known age-related metabolomic differences (Zhao et al., 2022; Helfand et al., 2003). For day 14 sampling, adult females were collected directly from vials, after which remaining flies were transferred to cages (∼1,000 individuals per population) and maintained until day 40, when the second sampling was performed. For each replicate population, samples were collected at three pool sizes (5, 50, and 100 individuals), selected to span the range commonly used in metabolomic studies and to maximize potential differences in detection (3 replicates × 2 genetic backgrounds × 3 pool sizes × 2 ages = 36 samples).

For Experiment 2, to assess how pool size interacts with replication and diet treatment, replicate ORWT populations were maintained under standard (STD) or high-sugar (HSD) diets (8 replicates per diet). Adult females were sampled 1–2 days post-eclosion for each replicate, standardizing developmental stage despite differences in development time between treatments. Samples were collected at pool sizes of 5, 50, and 100 individuals (2 diets × 8 replicates × 3 pool sizes = 48 samples).

For both experiments, flies were anesthetized on ice, transferred to a chilled ceramic plate, and sexed. Only females were used to eliminate sex-specific metabolic differences (Millington & Rideout, 2018). Individuals were pooled into the designated sample sizes, flash-frozen in liquid nitrogen, and stored at −80 °C until metabolomic analysis.

### Metabolomic Characterization and Data Processing

For untargeted metabolomics analyses, high resolution mass spectrometry (AB Sciex 5600 tripleTOF) was used. The whole fly samples were homogenized using a beads beater and extracted with a mixture of ice-cold methanol in water (1 mL, 4:1, v/v), incubated at −20 °C for 1 hour, then the homogenates centrifuged at 15,000 g at 4°C for 10 min. The 800 µL of supernatant was dried by speed-vac and then reconstituted with 200 µL of acetonitrile:water (1:1, v/v). The mixture was centrifuged at 15,000 g at 4°C for 10 min then supernatant was transferred into LC vials. Quality control (QC) sample was pooled from all samples to monitor analytical variability and to correct for any signal drift across batches of samples in untargeted metabolomics. Samples were analyzed by ultra-performance liquid chromatography coupled with electrospray quadrupole-time of flight (UPLC-QToF) mass spectrometry in the positive and negative ion modes with C18 and HILIC method using an Applied Biosystems Sciex 5600 instrument (Ref). Metabolites were identified based on accurate mass, MS/MS spectra, isotope pattern, and retention time using PeakView and MultiQuant software (AB Sciex) compared with metabolites in the facility’s metabolite IROA library (650 standards). When standards for comparison were not available, online databases, METLIN, HMDB, for metabolite identification were used (Choi et al., 2019).

Normalization was conducted independently for each experimental dataset. Relative concentration values for metabolites were log10-transformed to approximate a Gaussian distribution. Prior to normalization, metabolites with high technical variability were removed based on pooled quality control (QC) samples (coefficient of variation > 0.30). Data were then mean-centered within each analytical run using one of two approaches depending on the downstream analysis: (i) mean-centering by metabolite across samples to normalize metabolite-specific signal intensity differences while preserving variation among samples, or (ii) mean-centering within samples to normalize overall signal intensity differences while preserving relative metabolite composition within each sample.

### Metabolomic similarity and profile structure

To evaluate the effect of pool size on overall metabolomic composition, a common similarity analysis framework was applied to both experiments, with model terms adjusted to reflect the experimental design of each dataset. All analyses were performed separately for each LC–MS panel.

Following quality-control filtering, metabolite values were mean-centered within each sample across all metabolites in each panel. This normalization ensures that distances between samples reflect differences in metabolomic profile shape rather than overall differences in metabolite abundance.

Three complementary approaches were used to assess similarity. First, principal component analysis (PCA) was performed using the prcomp function in R to visualize clustering of samples by pool size in multivariate metabolomic space. Prior to PCA, metabolite values were mean-centered across samples so that principal components capture variation among samples rather than differences in baseline metabolite abundance. Additional scaling was not applied, as metabolites were already on a comparable scale following log-transformation and normalization (Hubert et al., 2025).

Second, pairwise Euclidean distances were calculated in the full metabolomic space between all pool-size pairs (5–50, 5–100, and 50–100). Distances were computed within each relevant biological grouping (see below), yielding one distance value per pool-size pair per replicate. These distances quantify the magnitude of metabolomic divergence between pool sizes, where a large 5–50 distance relative to a small 50–100 distance indicates that most profile change occurs at the first increase in pool size. Given the limited number of replicates, these comparisons are treated as descriptive and complement the formal statistical analyses.

Third, permutational multivariate analysis of variance (PERMANOVA) was performed using the adonis2 function from the *vegan* package (Oksanen et al., 2022) to test whether pool size explains a significant proportion of total metabolomic variance. Euclidean distance matrices were calculated from the normalized metabolite data for each panel. Models were fitted using marginal (Type III) tests so that the effect of pool size was evaluated after accounting for other experimental factors. Permutations (n = 999) were stratified within replicate populations using the how function from the *permute* package (Simpson, 2022) to account for non-independence of samples within replicates, and a fixed random seed was used for reproducibility. The primary output was R², representing the proportion of total variance explained by each factor.

Finally, multivariate dispersion was assessed using the betadisper function from the *vegan* package to evaluate the reproducibility of metabolomic profiles across pool sizes. Within each grouping, samples were assigned to pool-size categories, and the distance of each sample to its group centroid in multivariate space was calculated. A decrease in mean distance to centroid with increasing pool size indicates improved reproducibility. Differences in dispersion were tested using permutation tests (permutest, 999 permutations), with replicate population included as a blocking factor.

### Experiment-specific model structures

For **Experiment 1 (inbred vs. outbred populations)**, analyses were conducted within each strain × age × replicate combination for distance calculations and dispersion analyses. PERMANOVA models included pool size, strain, and age as fixed effects, allowing the contribution of pool size to be evaluated after accounting for genetic background and age-related differences. PCA was performed separately for each strain to avoid confounding pool-size effects with between-strain variation.

For **Experiment 2 (diet experiment)**, analyses were conducted within each diet × replicate combination for distance calculations and dispersion analyses. PERMANOVA models included pool size and diet as fixed effects, reflecting the absence of strain and age variation in this experiment. PCA was performed across all samples within each panel, as genetic background was held constant.

### Experiment 2: Effects of Pool Size and Replication on Signal Detection

#### Pool Size Analysis

To assess whether pool size affects the ability to detect genuine diet-related metabolomic changes, a per-metabolite signal detection analysis was performed using our four LC-MS panels. Input data were log-transformed metabolite values mean-centered across samples within each metabolite (as described above). For each metabolite in each panel, a linear model was fitted within each pool size:

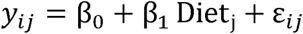

where *y_ij_* is the mean-centered, log-transformed abundance of metabolite *i* in sample *j*, Diet*_j_* is a two-level categorical predictor (HSD vs. STD), and ε*_ij_* is the residual error term. Models were fitted separately within each pool size (n = 5, n = 50, n = 100), using all 16 observations at that pool size (8 replicate populations × 2 diet groups). Running separate models per pool size ensure that results at each pool size reflect only the data collected at that pool size, allowing direct comparison of what a researcher would conclude from each pool size independently. To correct for multiple comparisons, p-values for the Diet coefficient were adjusted using the Benjamini-Hochberg false discovery rate (FDR) procedure applied within each pool size across all metabolites within a given panel. A metabolite was considered to have a significant diet effect at a given pool size if its FDR-adjusted p-value was below 0.05.

Results at n = 100 were treated as the ground truth reference, as this pool size provides the most stable and reproducible metabolomic profiles based on the similarity analysis. For each metabolite, the diet effect at n = 5 and n = 50 was classified relative to n = 100 using four categories: a true positive (TP) if the metabolite was significant at both pool sizes in the same direction; a false negative (FN) if the metabolite was significant at n = 100 but not at the smaller pool size; a false positive (FP) if the metabolite was significant at the smaller pool size but not at n = 100; and a true negative (TN) if the metabolite was not significant at either pool size. From these counts, two summary statistics were derived: sensitivity (TP / (TP + FN)), representing the proportion of genuinely diet-responsive metabolites recovered at a smaller pool size; and false discovery rate (FP / (FP + TP)), representing the proportion of detected effects that are not supported by the n = 100 reference.

Next, to assess whether pool size affects the reliability of individual effect size estimates, the diet coefficient estimated at n = 5 and n = 50 was compared against the corresponding estimate at n = 100 for all metabolites. Pearson correlation between effect sizes at each pool size and at n = 100 was computed separately for all metabolites and for the subset significant at n = 100, to assess whether the reliability of effect size estimates is consistent across the full metabolome.

Lastly, we quantified the sensitivity of individual metabolites to pool size by fitting a linear mixed-effects model across all pool sizes:

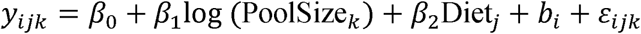

where *y_ijk_* is the mean-centered, log-transformed metabolite abundance, log (PoolSize*_k_*) is the natural log of pool size, and Diet*_j_* is a fixed effect (HSD vs. STD). The term *b_i_ ∼ N(0, a^2^_b_)* represents a random intercept for replicate population (pop_id). Pool size was modeled on the log scale because sampling effort increases nonlinearly across the discrete levels used (n = 5, 50, 100), and we were specifically interested in capturing diminishing returns in precision with increasing pool size. Models were fit in R using the “lme4” package, with t-statistics and p-values obtained using “lmertTest”. We quantified pool-size sensitivity as the absolute value of the t-statistic associated with log (PoolSize). Multiple testing correction was performed using the Benjamini–Hochberg procedure within each panel.

#### Downsampling & pool size

Replicate downsampling was used to assess the robustness of diet-associated metabolite signals to reduced biological replication. Analyses were performed separately for each normalized metabolomics dataset (C18 positive, C18 negative, HILIC positive, and HILIC negative) using data mean-centered by metabolite. Replicate populations were removed in a balanced manner across diets, with 0–5 replicate populations removed per diet and all possible combinations of removals evaluated at each level (Supplementary Table 2). For each downsampled dataset and each pool size (5, 50, and 100), metabolite abundance was modeled as a function of diet using simple linear models (metabolite ∼ Diet). Diet effect sizes, test statistics, and p-values were extracted for each metabolite, and p-values were adjusted for multiple testing using the Benjamini–Hochberg FDR procedure within each dataset × pool size × iteration combination. Metabolites with FDR < 0.05 were considered significant. Results were summarized across iterations to quantify changes in significant metabolite counts, true- and false-positive recovery relative to the reference condition (pool size = 100, full replication), concordance of diet effect sizes across pool sizes at full replication, and retention of significant metabolites across downsampling levels for metabolites grouped into high-, medium-, and low-effect bins based on reference absolute effect size.

#### Functional signal retention

To evaluate the retention of interpretable biological signal under replicate downsampling and varying pool size, metabolites were assigned to functional modules based on biochemical identity and established metabolic pathway context using curated classification rules informed by central metabolic processes. (Xia and Wishart, 2010; Kanehisa et al., 2012; Supplementary Table 3). Modules represented major metabolic processes, and metabolites not assigned to a defined module were excluded from module-level analyses. A reference set of diet-associated metabolites was defined from the full dataset (pool size = 100, maximum replication) using an FDR threshold of 0.05. Module-level signal preservation was quantified by calculating, for each downsampling condition, the proportion of reference metabolites that remained significant across iterations. Recovery frequency was first calculated at the metabolite level and then averaged within modules to obtain the percentage of metabolites recovered per module. Results were summarized across replicate downsampling levels and pool sizes and visualized as heatmaps to assess the robustness of functional metabolic signals.

#### Determinants of metabolite signal retention

To identify the factors underlying variation in metabolite signal retention across downsampling conditions, we modeled the probability that individual metabolites remained significant as a function of effect size, variability, pool size, and replication. For each metabolite within each dataset, pool size, and replicate downsampling condition, signal retention was quantified as the proportion of downsampling iterations in which the metabolite was detected as significant (FDR < 0.05).

Reference effect size for each metabolite was defined as the absolute value of the diet coefficient estimated under the reference condition (pool size = 100, maximum replication). To quantify variability in metabolite estimates across downsampling, we calculated the standard deviation of the diet coefficient across all iterations within each dataset × pool size × replicate condition. This measure captures the stability of effect size estimates under reduced sampling.

A linear mixed-effects model was used to evaluate the contribution of these factors to signal retention. The response variable was metabolite-level detection frequency, and fixed effects included reference effect size, variability (standard deviation of the diet coefficient), pool size (categorical: 5, 50, 100), and replicate number (3-8 replicates). Dataset and metabolite identity were included as random intercepts to account for non-independence among measurements from the same dataset and inherent differences among metabolites. Models were fit using the lme4 package in R.

## Results

### Experiment 1: Effects of Pool Size on Metabolomic Structure

#### Metabolomic similarity and profile structure

PCA revealed consistent separation of samples by pool size across all three LC-MS panels and both strains (Figure 1). In all panels, n=5 samples were consistently separated from n=50 and n=100 samples along PC1, which explained 46–62% of total variance, while n=50 and n=100 samples clustered more closely together. Separation by age was also visible along PC2, particularly in the HILIC panels, consistent with known age-related metabolomic changes.

**Figure 1.**
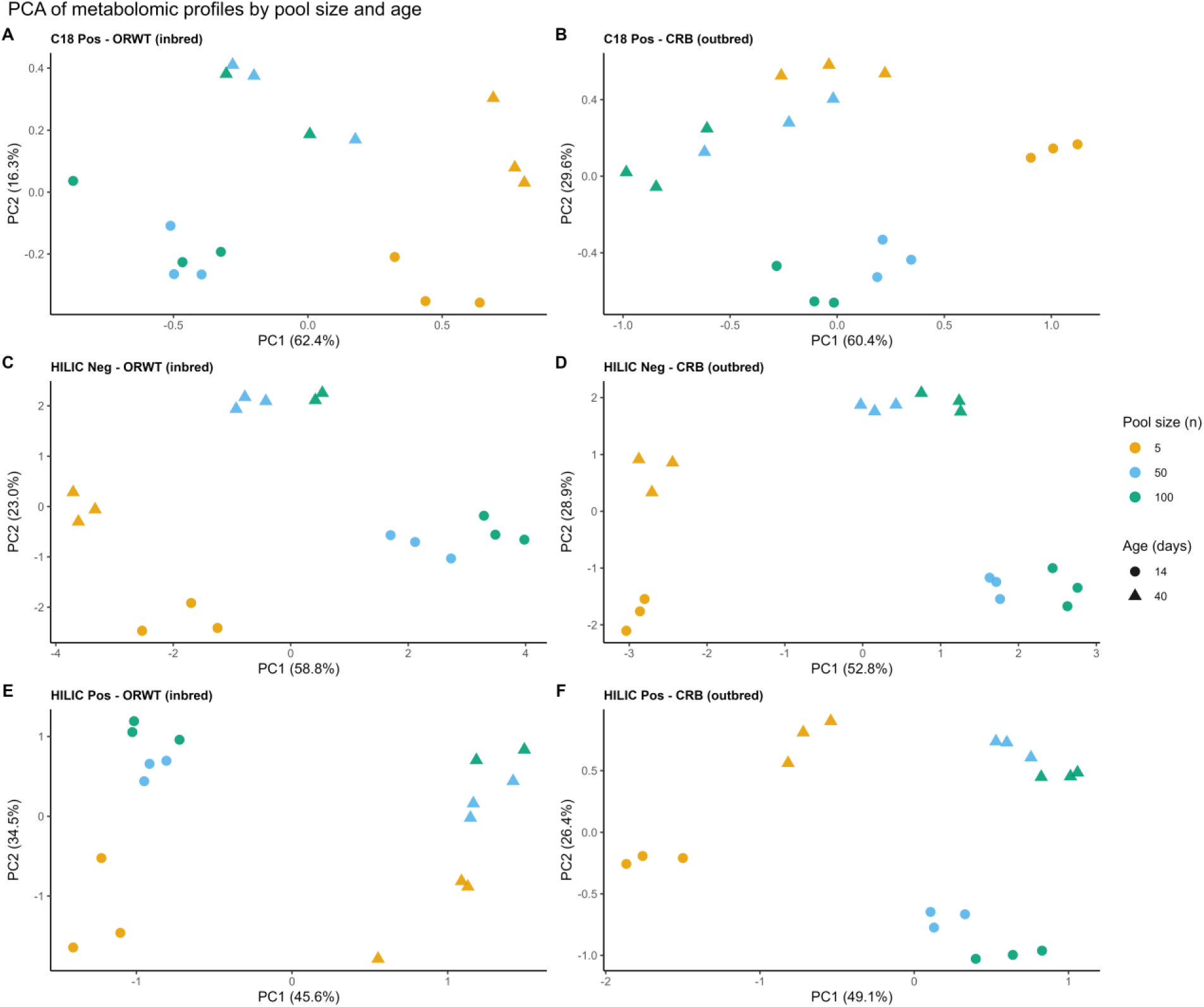
PCA of metabolomic profiles by pool size and age across three metabolite panels. Points represent individual samples colored by pool size and shaped by age. PCA was performed on mean-centered metabolite values without additional scaling. Variance explained by each PC is shown in parentheses on axis labels.

Pairwise Euclidean distances were consistent with the PCA, with n=5 profiles being more dissimilar to larger pools than the larger pools were to each other (Figure 2, Supplementary Table 4). The effect was strongest in the HILIC panels, where 5–100 distances were 2.3–3.9 times larger than 50–100 distances across all strains and ages, with HILIC negative showing the greatest absolute distances reflecting its larger metabolite count. In C18 positive the same directional pattern held in most conditions, though the difference between pairs was less pronounced (ratio range 1.6–2.8×), with the notable exception of older CRB flies where 5–50 and 50–100 distances were nearly equivalent (0.66 and 0.64 respectively). Across all panels, 5–50 distances were consistently intermediate, indicating that most profile divergence occurs at the first pooling step, with comparatively little additional change from n=50 to n=100.

**Figure 2.**
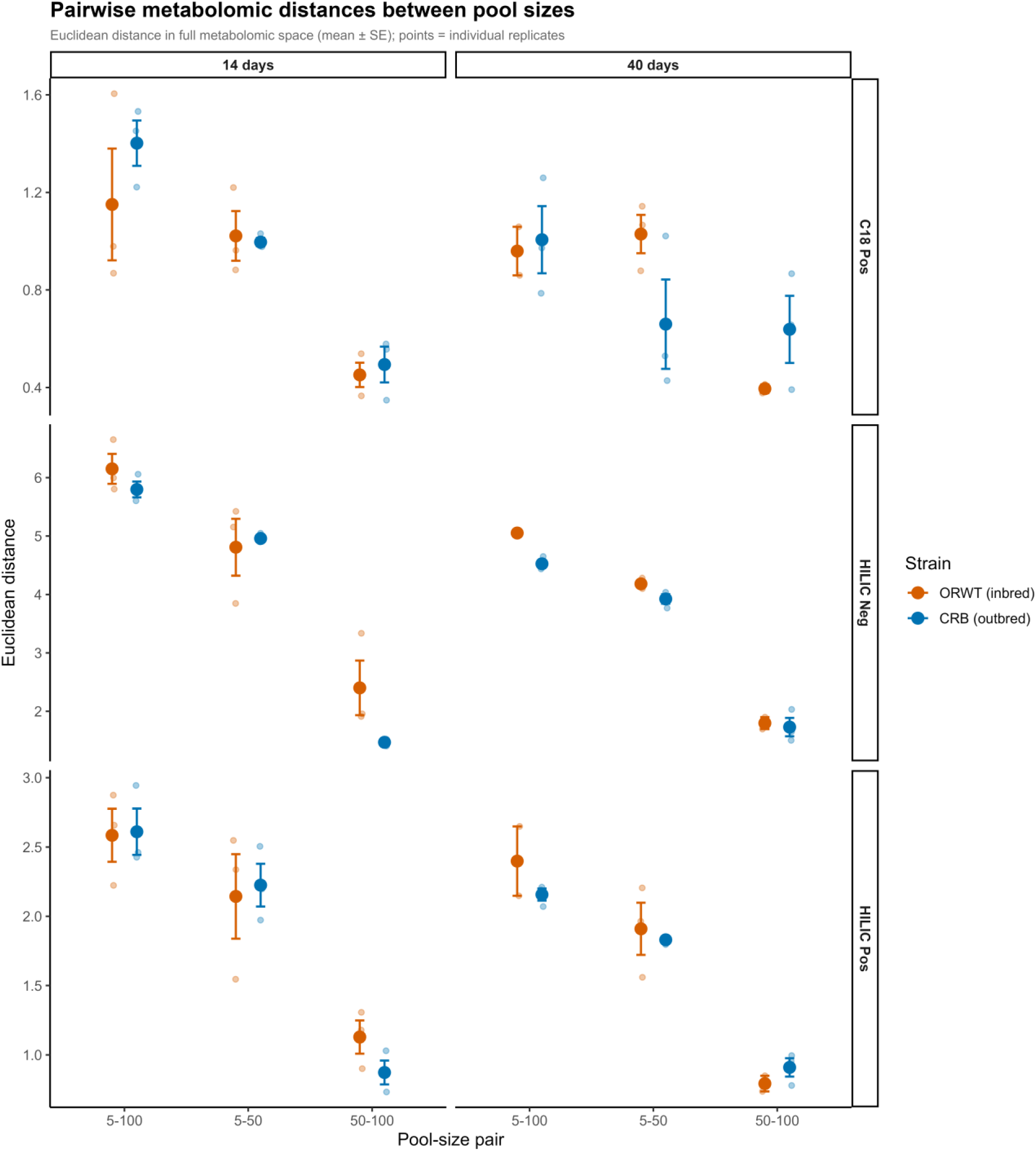
Pairwise Euclidean distances between pool sizes in the full metabolomic space. Mean distance (± SE) between all pool-size pairs (5–100, 5–50, 50–100) is shown for each metabolite panel, coloured by strain. Individual replicate values are shown as transparent points.

PERMANOVA confirmed that pool size explained a statistically significant proportion of metabolomic variance in all three panels, though the magnitude varied considerably (Figure 3A). The effect was largest in HILIC negative (R² = 0.333, p = 0.001) and HILIC positive (R² = 0.151, p = 0.001), and notably smaller in C18 positive (R² = 0.057, p = 0.001), where strain was the overwhelmingly dominant source of variation (R² = 0.695, p = 0.001). Strain was a significant predictor across all panels (R² = 0.262–0.695, all p = 0.001), as was age (R² = 0.029–0.189, all p = 0.001), with the largest age effect in HILIC negative. The weaker pool size signal in C18 positive likely reflects the fact that strain differences account for so much of the total variance in that panel that the pool size effect, while present, represents a smaller relative fraction. Importantly, pool size remained significant after marginal adjustment for both strain and age in all panels, confirming that its effect on metabolomic composition is independent of biological group differences.

**Figure 3.**
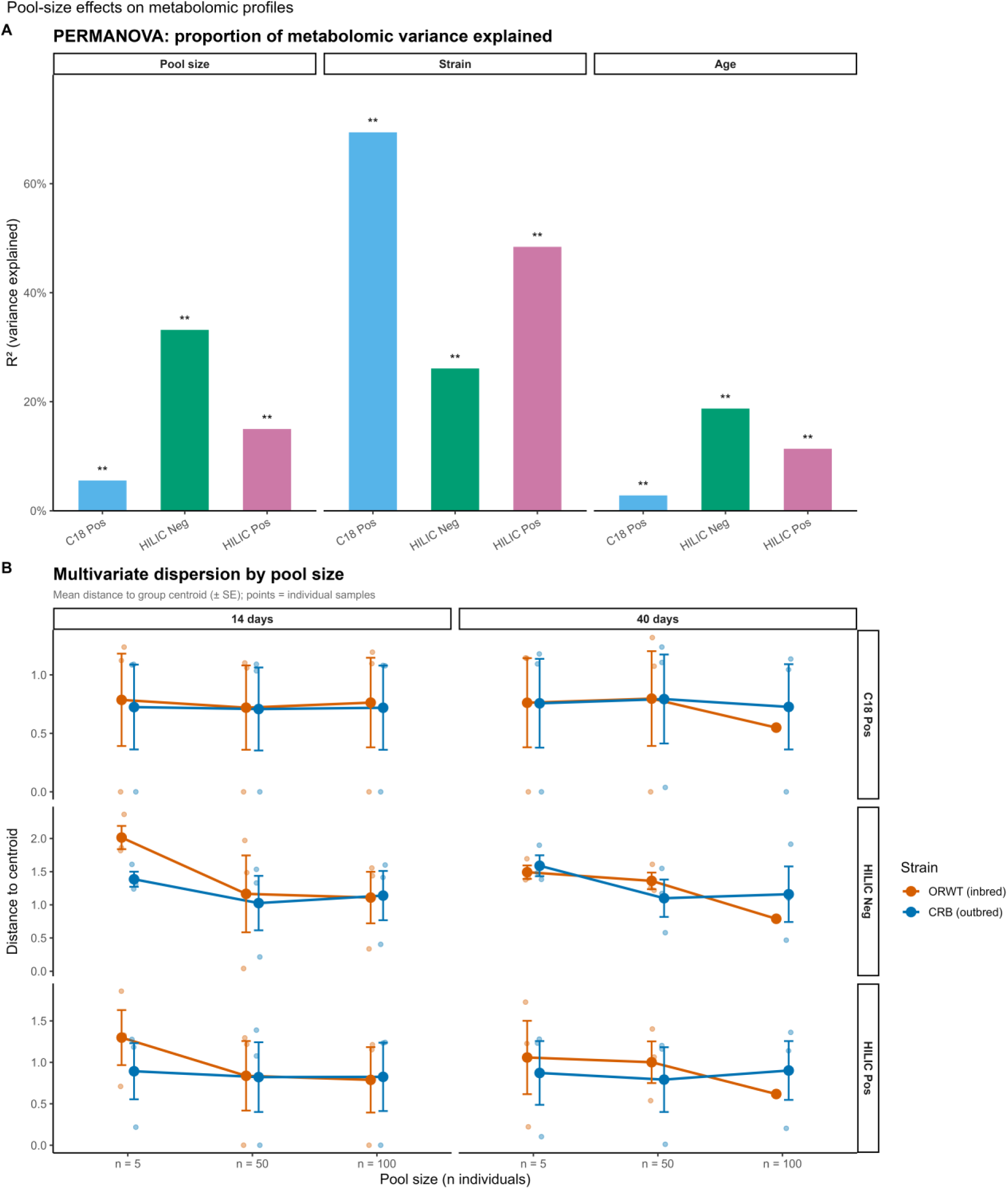
Pool-size effects on metabolomic profiles. (A) Proportion of total metabolomic variance explained by pool size, strain, and age for each panel based on PERMANOVA results. (B) Multivariate dispersion by pool size, shown as mean distance to group centroid (± SE) within each strain × age combination. Individual sample values are shown as transparent points.

Betadisper analysis showed that the decline in multivariate dispersion with increasing pool size was clearest in HILIC negative, consistent with the largest pool size R² in that panel, and more variable elsewhere (Figure 3B). In HILIC negative, mean dispersion at n=5 ranged from 1.39 to 2.01 across strain and age subgroups, falling to 1.03–1.36 at n=50 and 0.79–1.16 at n=100, indicating a consistent improvement in reproducibility with larger pool sizes. In HILIC positive, a directional decline was present in the ORWT strain (n=5: 1.06–1.30; n=50: 0.84–1.00) but was less consistent in CRB, where dispersion was similar across all pool sizes (0.82–0.89). In C18 positive, dispersion was low and stable across all pool sizes and subgroups (0.55–0.80), with no directional pattern, in keeping with the weak pool size effect seen in PERMANOVA. Together, these results indicate that pooling at least 50 individuals improves profile reproducibility most clearly in HILIC negative, with a strain-dependent pattern in HILIC positive and negligible pool-size effects on reproducibility in C18 positive.

### Experiment 2: Effects of Pool Size and Replication on Signal Detection

#### Metabolomic similarity and profile structure

PCA showed consistent separation by pool size across all metabolite panels, with n = 5 samples clearly separated from n = 50 and n = 100, which clustered more closely (Figure 4), mirroring the pattern observed in Experiment 1. PC1 explained 51–64% of total variance across panels. Along PC2, samples showed separation by diet, most clearly in the C18 panels and to a lesser extent in HILIC negative, while HILIC positive exhibited greater scatter and less distinct clustering. This likely reflects the smaller number of metabolites in this panel (n = 18) relative to the others.

**Figure 4.**
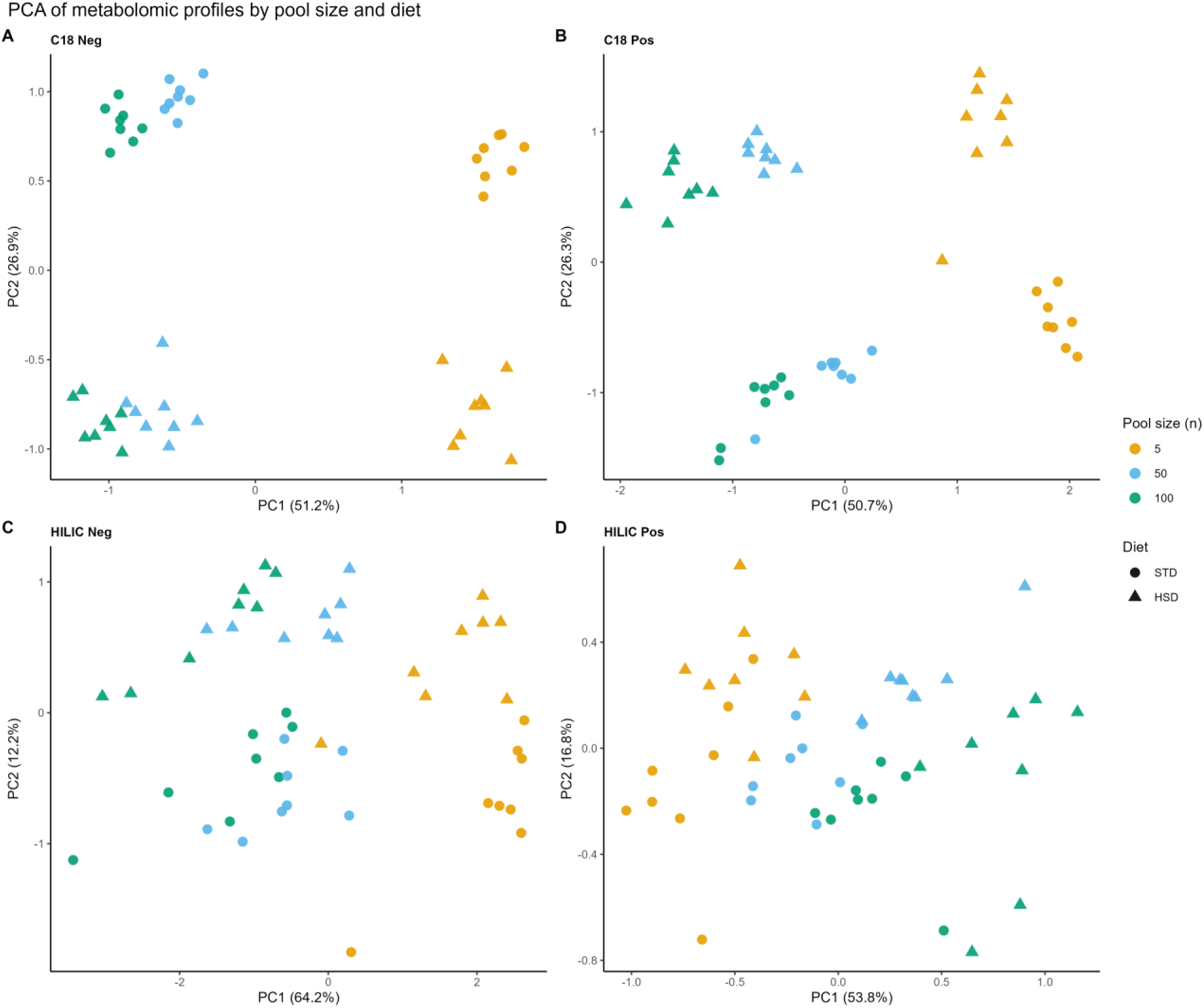
PCA of metabolomic profiles coded by pool size and diet across four metabolite panels. Points represent individual samples colored by pool size and shaped by diet. PCA was performed on mean-centered metabolite values without additional scaling. Variance explained by each PC is shown in parentheses on axis labels.

Pairwise Euclidean distances confirmed this pattern, with n = 5 profiles substantially more dissimilar to larger pool sizes than n = 50 and n = 100 were to each other (Figure 5, Supplementary Table 5). Across all panels, mean 5–100 distances were approximately 1.8–2.8 times larger than 50–100 distances, with 5–50 distances consistently intermediate. These patterns were consistent across diets, indicating that most profile divergence occurs at the first increase in pool size.

**Figure 5.**
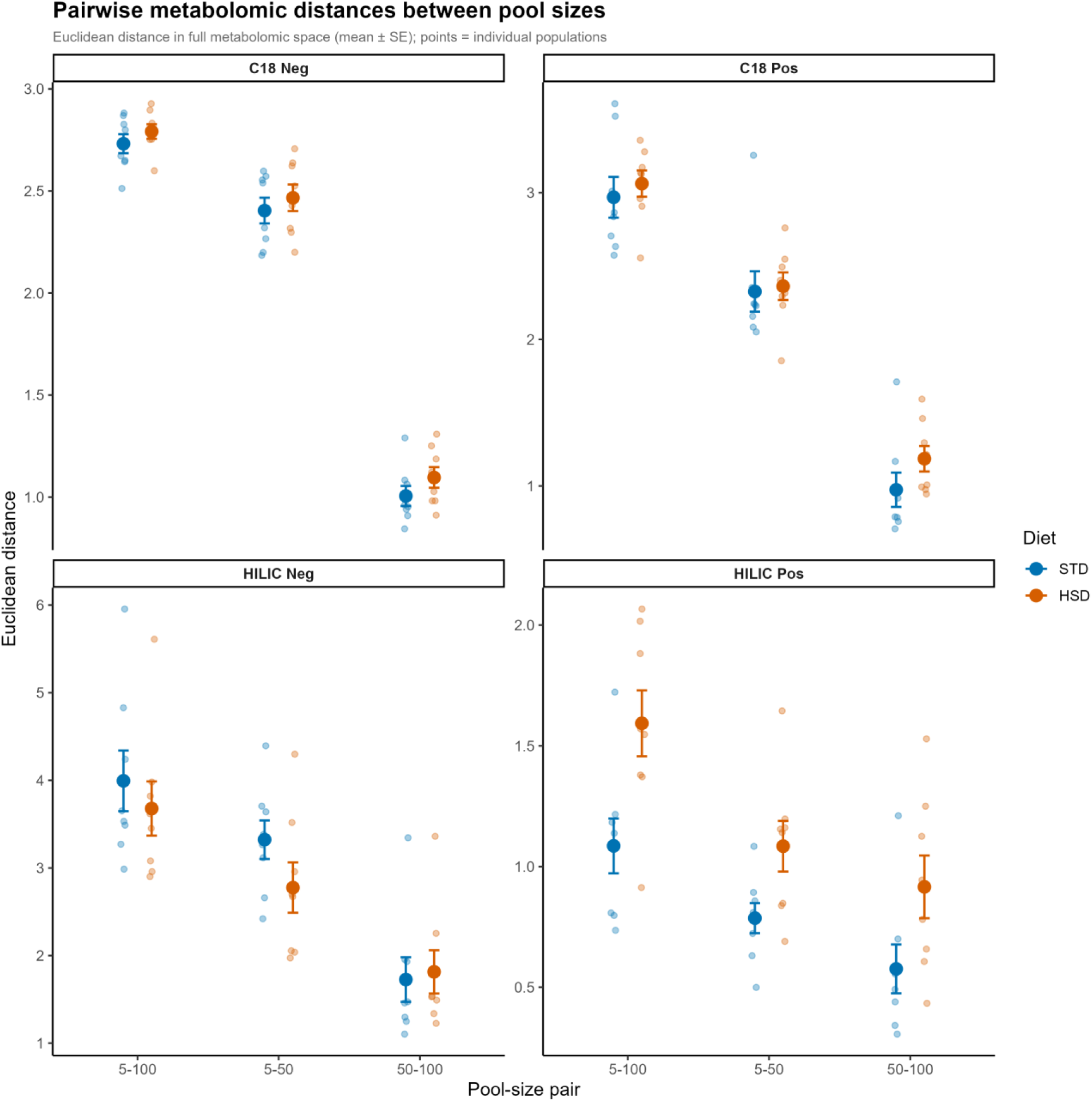
Pairwise Euclidean distances between pool sizes in the full metabolomic space. Mean distance (± SE) between all pool-size pairs (5–100, 5–50, 50–100) is shown for each metabolite panel, colored by diet.Individual replicate values are shown as transparent points.

PERMANOVA further showed that pool size explained a large proportion of metabolomic variance in all panels (Figure 6A), accounting for 39–54% of total variance depending on the panel. Diet was also a significant predictor, explaining 11–27% of variance, with stronger effects in the C18 panels relative to the HILIC panels. The persistence of a strong pool size effect after accounting for diet indicates that pooling influences metabolomic composition independently of treatment effects.

**Figure 6.**
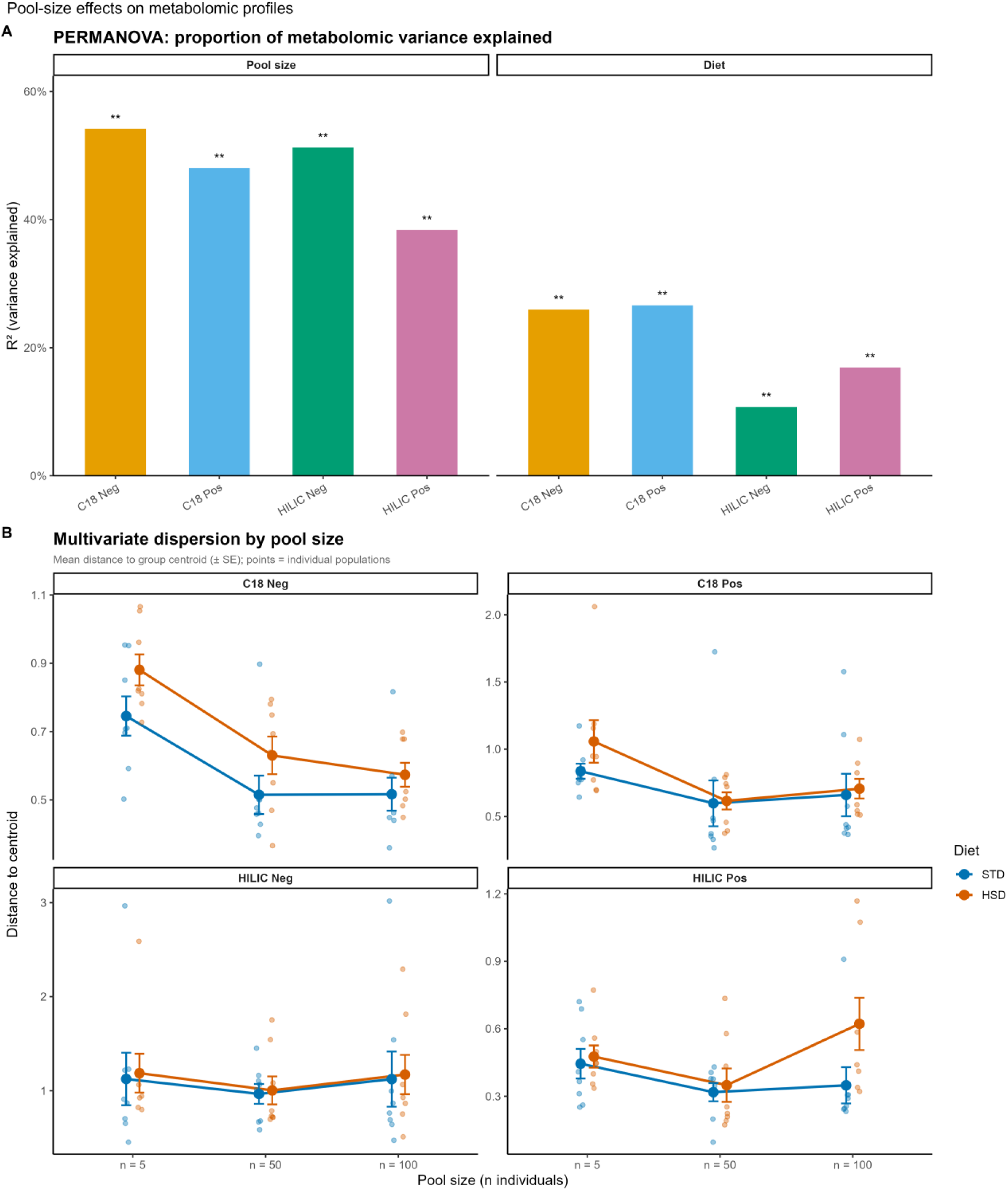
Pool-size effects on metabolomic profiles. (A) Proportion of total metabolomic variance explained by pool size and diet for each panel based on PERMANOVA results.(B) Multivariate dispersion by pool size, shown as mean distance to group centroid (± SE) within each diet group.

Betadisper analysis revealed that improvements in reproducibility with increasing pool size were most pronounced in the C18 panels (Figure 6B). In both C18 panels, dispersion declined substantially from n = 5 to n = 50, with little additional improvement at n = 100, indicating diminishing returns beyond intermediate pool sizes. In contrast, the HILIC panels showed weaker and less consistent patterns, with HILIC negative exhibiting relatively stable dispersion across pool sizes and HILIC positive showing variable behavior, including an increase in dispersion at n = 100 under the HSD treatment. This latter pattern should be interpreted cautiously given the small number of metabolites in this panel.

Together, these results confirm that pool size has a strong and nonlinear effect on metabolomic profile structure, with the largest changes occurring between n = 5 and n = 50. Compared to Experiment 1, the magnitude of the pool size effect was substantially larger in this dataset, and the presence of a strong diet signal further demonstrates that pooling effects persist even in the presence of pronounced biological differences.

#### Detection of diet-associated metabolites across pool sizes

To assess the practical consequences of pool size choice on biological inference, we examined whether pool size affects the ability to detect diet-related changes in individual metabolite levels. For each metabolite and pool size, a linear model was fitted to test the effect of diet, and results at n = 100 were treated as the ground truth reference. Sensitivity was substantially lower at n = 5 than at n = 50 across all panels (Figure 7A, Supplementary Table 6D). At n = 5, sensitivity ranged from 50% in both HILIC negative and HILIC positive to 73% in C18 positive, meaning that between 27% and 50% of real diet effects would be missed depending on the panel. At n = 50, sensitivity improved substantially across all panels, reaching 89% in C18 negative, 90% in C18 positive, 75% in HILIC negative, and 100% in HILIC positive, though the HILIC positive result should be interpreted cautiously given the small number of metabolites in that panel (18 metabolites) and the limited number of significant effects at n = 100 (n = 10).

**Figure 7.**
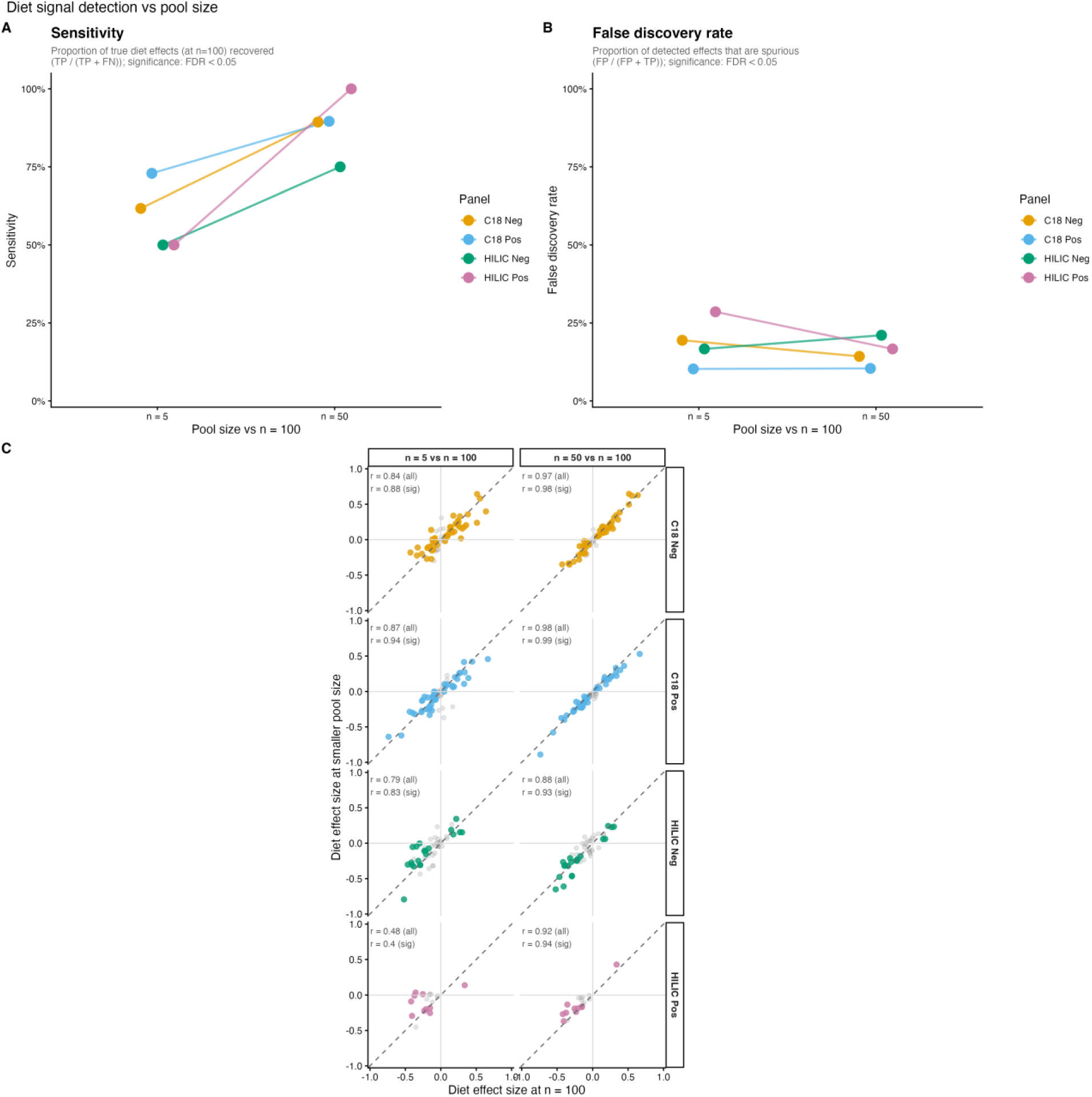
Diet signal detection vs pool size. (A) Sensitivity — proportion of true diet effects (significant at n = 100, FDR < 0.05) recovered at n = 5 or n = 50. (B) False discovery rate — proportion of detected effects not supported at n = 100. Each point is one LC-MS panel; lines connect the two pool-size comparisons. (C) Diet effect sizes at smaller pool sizes versus n = 100 for all metabolites. Colored points are significant at n = 100 (FDR < 0.05); grey points are not. Pearson r shown for all metabolites and the significant subset.

The false discovery rate (FDR) was low and broadly consistent across panels at both pool sizes (Figure 7B, Supplementary Table 6B). In C18 negative, FDR was 19% at n = 5 and 14% at n = 50; in C18 positive, FDR was 10% at n = 5 and remained similar at n = 50 (10%). HILIC negative showed FDR of 17% at n = 5 rising modestly to 21% at n = 50. HILIC positive showed FDR of 29% at n = 5 dropping to 17% at n = 50, though as noted above the absolute counts are small.

Effect size attenuation analysis confirmed that pool size does not systematically bias the direction or magnitude of diet effect estimates, but does introduce substantial noise at n = 5 (Figure 7C, Supplementary Table 6C). Across all metabolites, Pearson correlations between diet effect sizes at smaller pool sizes and at n = 100 were consistently high at n = 50 (r = 0.92–0.98 across panels), with points clustering tightly around the 1:1 line. At n = 5, correlations were lower and more variable across panels (r = 0.48–0.88), with HILIC positive showing the greatest scatter (r = 0.48) and C18 positive the highest reliability (r = 0.87). Notably, metabolites with the largest diet effect sizes at n = 100 showed the greatest consistency across pool sizes, clustering close to the 1:1 line even at n = 5. The increased scatter at n = 5 was concentrated among metabolites with modest effect sizes, suggesting that pools of n = 5 are adequate for detecting the strongest diet signals but produce unreliable estimates for moderate and small effects. Correlations were similar whether computed across all metabolites or restricted to those significant at n = 100, indicating that this reliability pattern is representative of the metabolome as a whole rather than an artefact of pre-selecting metabolites with large effects.

The supplementary sensitivity analysis confirmed that pool size has a strong and nearly universal effect on individual metabolite values (Supplementary Figure 1). Across all panels, the majority of metabolites showed |t|-values well above the approximate significance threshold of 2 for the log(pool size) term, and almost all metabolites showed a positive relationship with pool size. This directional consistency indicates that pool size introduces a systematic upward shift in measured metabolite levels rather than random noise, likely reflecting the averaging of individual stochasticity in small pools. The C18 panels showed notably higher pool-size sensitivity than the HILIC panels, consistent with the stronger pool size effects in these panels observed in the similarity analysis.

#### Impact of replicate downsampling on signal detection

To evaluate the robustness of diet-associated metabolite signals, we systematically reduced replicate number across pool sizes and quantified the retention of significant metabolites relative to a reference condition (pool size = 100, full replication) (Supplementary Table 7). Across all datasets, the number of diet-significant metabolites declined with decreasing replicate number, with more pronounced losses observed at smaller pool sizes (Figure 8). At the largest pool size (100 individuals), a high proportion of the reference signal was retained at moderate replication, with gradual declines as replicate number decreased. In contrast, smaller pool sizes accelerated signal loss, particularly at pool size = 5, where the number of detected metabolites declined sharply even when several replicates remained.

**Figure 8.**
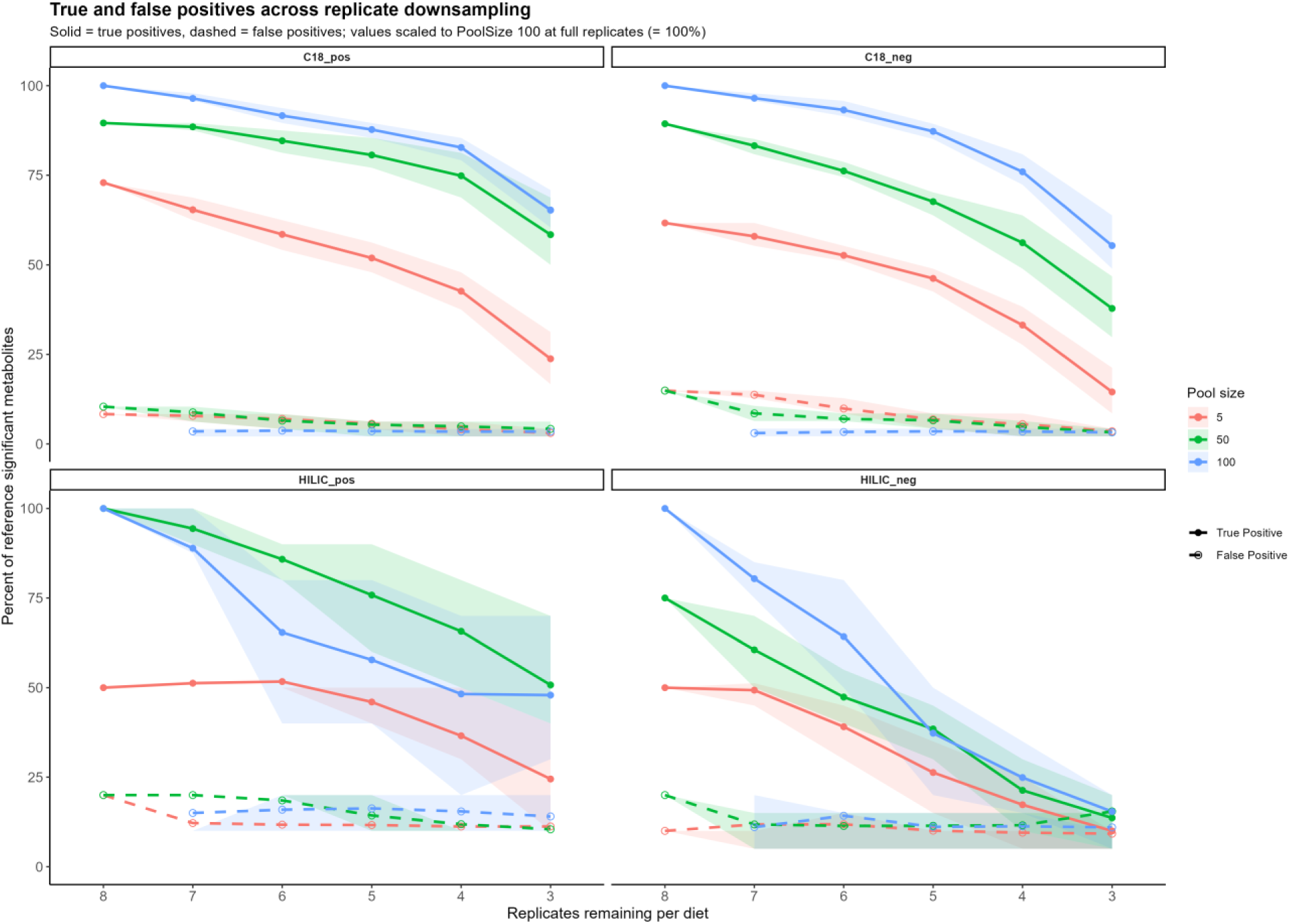
Retention of true and false positive metabolite detection across replicate and pool-size downsampling. The proportion of diet-associated metabolites identified under downsampling was expressed as a percentage of the reference set defined at PoolSize = 100 with full replicates (8 per diet; 100%). True positives (solid lines) represent metabolites that were significant (FDR < 0.05) in both the downsampled and reference datasets, while false positives (dashed lines) were significant only in the downsampled condition. Lines show the mean percentage across all combinations of replicate removal, and shaded ribbons indicate the interquartile range (25th–75th percentile), reflecting variability across downsampling iterations. Across all metabolite panels, reductions in replicate number and pool size led to a progressive loss of true positives, while false positives remained comparatively low, indicating reduced statistical power rather than systematic inflation of spurious detections under downsampling.

Classification of significant metabolites relative to the reference condition revealed that reductions in replicate number primarily resulted in the loss of true positives rather than an increase in false positives. Across all datasets and pool sizes, the proportion of true positives decreased steadily with replicate removal, while false positives remained comparatively low and showed only modest increases at low replication levels. These patterns indicate that, similar to our findings of reduced pool size, reduced replication primarily limits statistical power rather than inflating spurious detections.

#### Effect size retention across pool sizes under replicate downsampling

To determine whether signal stability depended on the magnitude of the underlying biological effect, metabolites were grouped into high-, medium-, and low-effect bins based on absolute diet effect size under the reference condition. Signal retention differed markedly across these categories (Figure 9). High-effect metabolites remained detectable across a wide range of downsampling conditions, whereas medium-effect metabolites showed more gradual declines and low-effect metabolites were rapidly lost even with modest reductions in replication.

**Figure 9.**
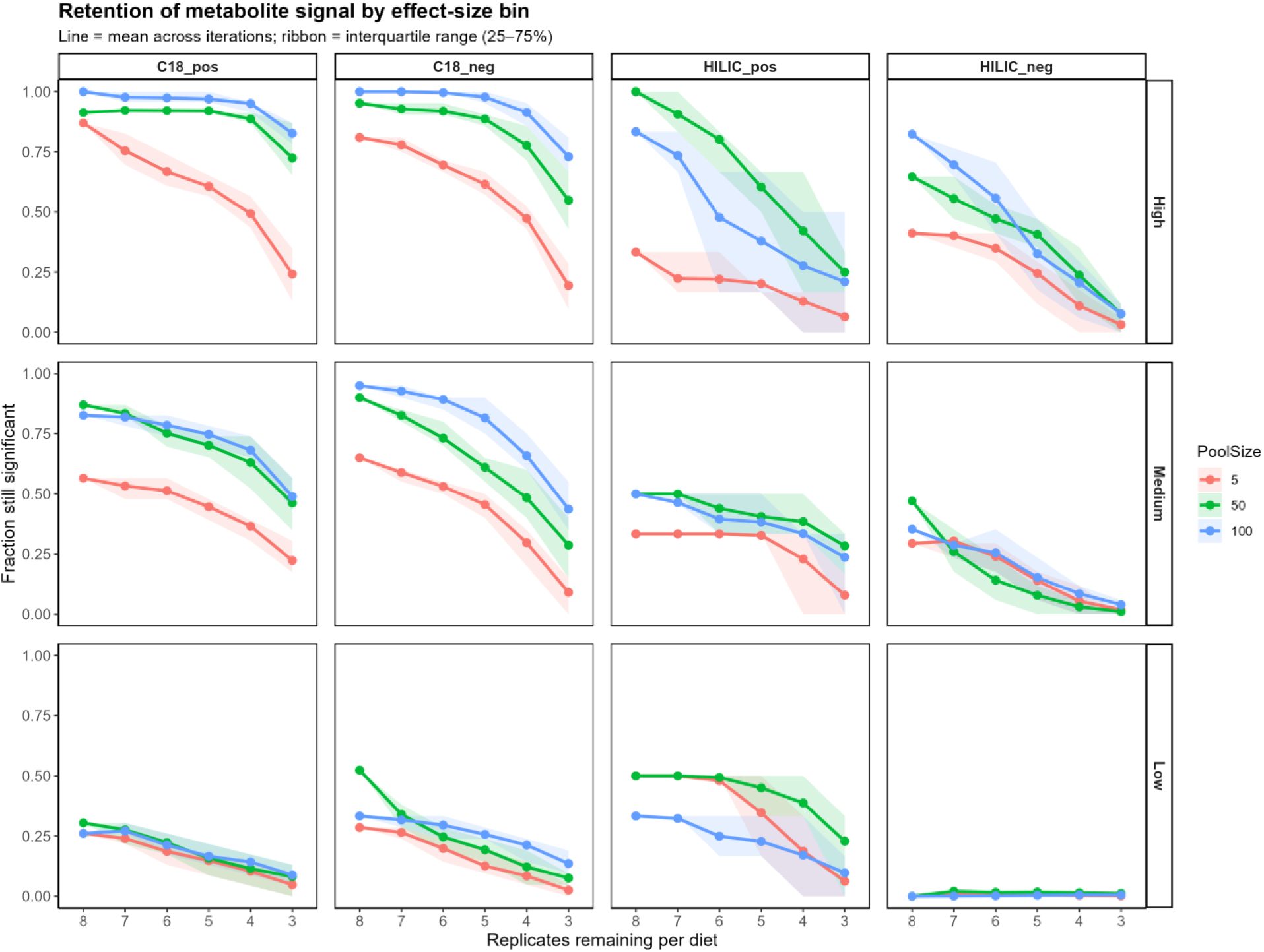
Retention of metabolite signal across replicate and pool-size downsampling, stratified by effect-size bin. Metabolites were grouped into high, medium, and low effect-size bins based on tertiles of absolute diet effect sizes estimated from the full dataset (PoolSize = 100, full replicates), calculated separately within each metabolite panel. For each bin, the proportion of metabolites remaining significant (FDR < 0.05) was evaluated across all combinations of replicate downsampling and pool sizes (5, 50, and 100). Lines represent the mean fraction of metabolites remaining significant across all downsampling iterations, and shaded ribbons indicate the interquartile range (25th–75th percentile), reflecting sensitivity to which replicate populations were removed. Across all panels, metabolites with larger effect sizes exhibited greater robustness to reductions in replicate number and pool size, whereas low-effect metabolites rapidly lost significance under downsampling.

While all datasets exhibited a consistent directional trend of decreasing detection with replicate loss, the rate of signal decline varied substantially across datasets. In some datasets, high- and medium-effect metabolites were retained across a broader range of downsampling levels, whereas in others, detection declined more rapidly across all effect-size categories. These differences indicate that, in addition to effect size, dataset-specific properties influence the robustness of metabolite detection under reduced replication.

In addition to effects on statistical significance, replicate downsampling also influenced the magnitude and consistency of estimated diet effect sizes. At full replication, effect size estimates were highly concordant across pool sizes, with strong agreement between estimates from pool size of 50 relative to the reference pool size of 100. However, as replicate number decreased, effect size estimates became increasingly variable, with this effect strongly modulated by pool size. Smaller pool sizes, particularly pool size = 5, exhibited substantially greater variability and more rapid degradation of effect size estimates compared to larger pool sizes.

Consistent with this pattern, the rate of signal loss under downsampling differed markedly across pool sizes. Larger pool sizes retained diet-associated signals across a broader range of replicate reductions, whereas smaller pool sizes showed accelerated loss of both effect size stability and statistical detection. This effect was most pronounced for metabolites with smaller effect sizes, which were rapidly lost under conditions of both low replication and small pool size.

Overall, these results indicate that both biological replication and pool size jointly determine the robustness of diet-associated metabolite signals. These results demonstrate that pool size and replication are not interchangeable, but instead interact to determine the reliability of metabolomic inference. Larger pool sizes reduce variability and stabilize effect size estimates, thereby slowing the rate at which signals are lost under reduced replication, while increased replication improves detection across all conditions. Together, these factors jointly determine the effective signal-to-noise ratio and, consequently, the ability to detect consistent biological effects.

#### Functional signal retention across sampling conditions

Diet significantly altered metabolite profiles across multiple functional modules in the reference dataset (pool size = 100, maximum replication). The strongest responses were observed in lipid-associated pathways, including fatty acid oxidation and fatty acid overflow, as well as central carbon metabolism and one-carbon metabolism (Figure 10A). These coordinated changes are consistent with increased carbon flux and lipid remodeling under high-sugar conditions, indicating system-wide metabolic reprogramming.

**Figure 10.**
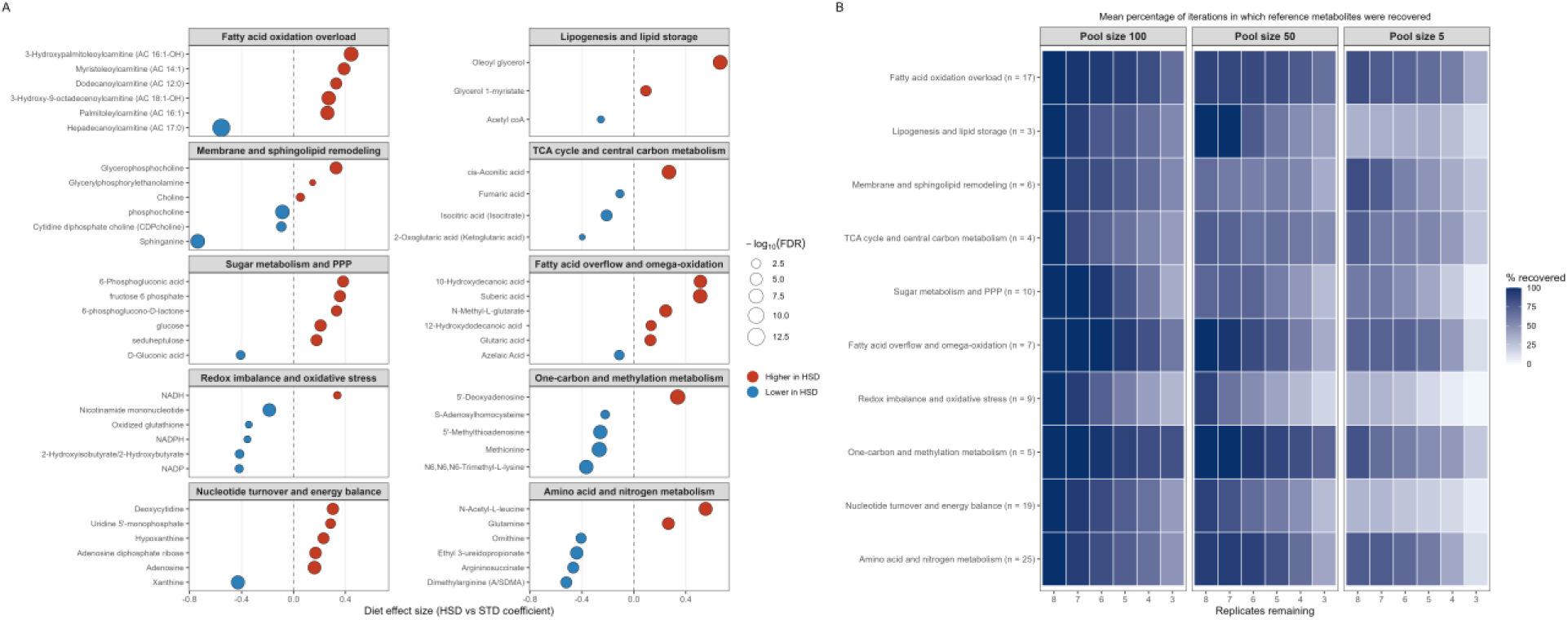
Metabolic responses to diet and preservation of signals under downsampling. (A) Top diet-responsive metabolites from the reference dataset (pool size = 100, maximum replication) grouped by functional module. Points represent individual metabolites, with position indicating effect size (HSD vs STD), size reflecting statistical significance (−log10 FDR), and color indicating direction of change. (B) Heatmaps showing the percentage of reference metabolites recovered (FDR < 0.05) across decreasing replicate number and pool sizes (100, 50, and 5 individuals; panels). Values represent the mean recovery across iterations. Metabolites are grouped into functional modules (module sizes indicated on the y-axis). Recovery declines with reduced replication and smaller pool sizes, with lipid and one-carbon metabolism remaining relatively robust, whereas redox and nucleotide metabolism show rapid loss of signal.

To evaluate the robustness of these signals to reduced sampling effort, we quantified the retention of reference metabolites across replicate downsampling and pool size reduction (Figure 10B). Across all modules, the proportion of recovered metabolites declined with decreasing replicate number, with additional losses observed at smaller pool sizes. At the largest pool size (100 individuals), most modules retained a high proportion of the reference signal at moderate replication, with gradual declines as replicate number decreased. In contrast, smaller pool sizes accelerated signal loss, particularly at pool size = 5, where recovery declined sharply even at higher replicate numbers.

The extent of signal loss varied across metabolic modules. Lipid-associated pathways, including fatty acid oxidation and fatty acid overflow, as well as one-carbon metabolism, showed relatively high retention across conditions, remaining detectable under reduced replication and smaller pool sizes. In contrast, redox metabolism and nucleotide turnover exhibited rapid declines in recovery, indicating greater sensitivity to reduced sampling effort. Central carbon metabolism and amino acid metabolism showed intermediate behavior, with moderate decreases in retention as replication and pool size were reduced.

Together, these results again demonstrate that both replicate number and pool size influence the detection of metabolite-level signals. Importantly, the robustness of signal retention differed across metabolic pathways, indicating that some biological processes are more resilient to experimental downsampling than others.

#### Determinants of metabolite signal retention

To identify the factors underlying variation in metabolite signal retention across downsampling conditions, we modeled detection frequency as a function of effect size, variability, pool size, and replication using a linear mixed-effects model (Table 1). Across all datasets, retention was strongly positively associated with the magnitude of the underlying diet effect (t = 12.45), indicating that metabolites with larger effect sizes are substantially more robust to replicate downsampling.

**Table 1.**
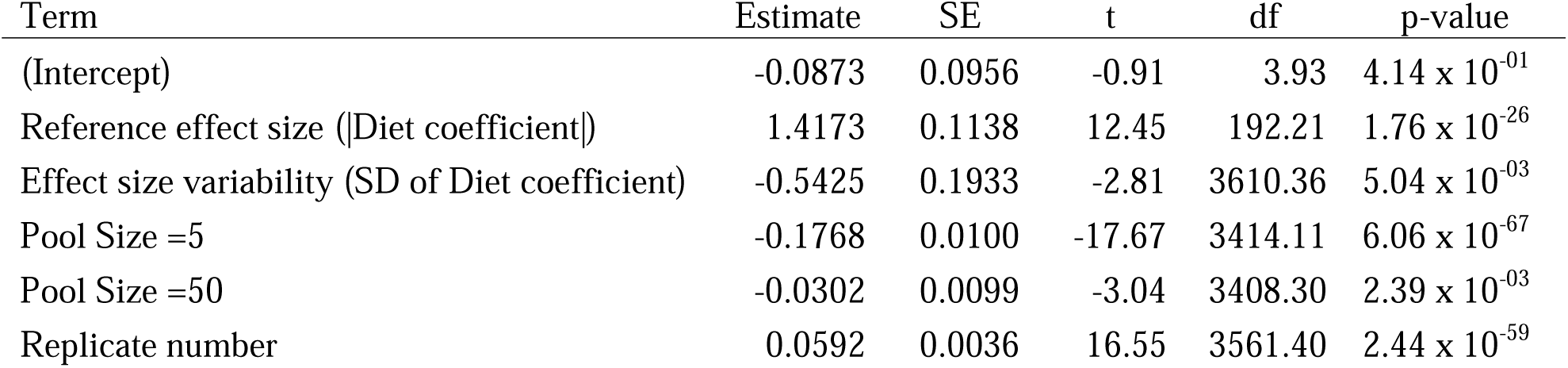
Determinants of metabolite signal retention under replicate downsampling. Linear mixed-effects model evaluating the effects of reference effect size, variability (standard deviation of the diet coefficient), pool size, and replicate number on metabolite detection frequency. Dataset and metabolite identity were included as random effects.

In contrast, variability in effect size estimates across iterations was negatively associated with retention (t = −2.81), demonstrating that metabolites with less stable estimates are more likely to be lost under reduced sampling. This relationship provides a mechanistic explanation for differences in signal robustness, linking reduced detection to increased variability rather than systematic bias.

Both pool size and replication had strong independent effects on detection. Pool size = 5 showed a substantial reduction in retention relative to pool size = 100 (t = −17.67), whereas pool size = 50 showed a smaller but still significant reduction (t = −3.04). These results are consistent with a nonlinear improvement in detection between 5 and 50 individuals and diminishing returns at larger pool sizes. Increasing replicate number strongly improved detection (t = 16.55), reinforcing the importance of biological replication across all conditions.

Together, these results demonstrate that metabolite detection is governed by a balance between biological signal strength and measurement variability, with pool size and replication jointly modulating the ability to detect consistent effects. Differences in signal robustness across datasets are therefore explained not only by experimental design, but also by underlying variation in effect size and stability among metabolites.

## Discussion

Pooling is a necessary but highly variable component of metabolomics experimental design, particularly in small organisms such as *Drosophila melanogaster* (Zhao et al., 2022; Hubert et al., 2025; Vecchié et al., 2026). Despite its widespread use, the consequences of pool size for metabolomic measurement and biological inference remain poorly defined. Here, using two complementary experimental designs, we show that pool size and biological replication jointly determine both the structure of metabolomic profiles and the ability to detect biologically meaningful signals. Importantly, these effects are nonlinear and dataset-dependent, with the most substantial differences occurring between very small pools (n = 5) and larger pools (n ≥ 50).

### Pool size fundamentally alters metabolomic profile structure

In Experiment 1, we found that pool size fundamentally alters the structure of metabolomic profiles. Across all panels, samples pooled at n = 5 were consistently separated from those pooled at n = 50 and n = 100, with much smaller differences observed between the two larger pool sizes. Pairwise distance analyses further showed that most of the divergence in metabolomic profiles occurs at the first increase in pool size, from 5 to 50 individuals, with relatively little additional change from 50 to 100. These results indicate that small pools capture a substantially different representation of the metabolome than larger pools, likely reflecting increased influence of individual-level stochasticity and heterogeneity (Kendziorski et al., 2005). In contrast, larger pools appear to more reliably capture the average physiological state of the population.

Importantly, the inclusion of both inbred and outbred populations provides insight into how genetic background interacts with metabolomic variation. As expected, strain explained a substantial proportion of total variance across all panels, indicating strong underlying genetic effects on metabolomic profiles. However, the impact of pool size on profile structure and reproducibility did not differ consistently between inbred and outbred populations across datasets. While outbred populations are expected to exhibit greater individual-level variability, which could amplify the effects of pooling, our results suggest that the benefits of increased pool size are not determined solely by genetic variation (Mackay et al., 2012). Instead, pooling effects appear to depend on additional dataset-specific factors, including metabolite composition and measurement characteristics.

The magnitude of the pool size effect, however, varied substantially across datasets. Pool size explained a large proportion of variance in the HILIC panels, particularly HILIC negative, but a much smaller proportion in the C18 positive panel, where strain differences dominated the overall signal. Similarly, improvements in reproducibility with increasing pool size were most evident in HILIC negative, more variable in HILIC positive, and minimal in C18 positive. These results demonstrate that while pooling can improve the stability of metabolomic measurements, its impact is not uniform and depends on the underlying structure of the dataset, including the types of metabolites measured and the relative contribution of biological versus technical variation.

In Experiment 2, we directly evaluated how pool size affects biological inference by quantifying the detection of diet-associated metabolite changes. Smaller pool sizes, particularly n = 5, showed substantially reduced sensitivity, missing a large fraction of diet-responsive metabolites identified at n = 100. In contrast, false discovery rates remained relatively low and stable across pool sizes, indicating that reduced pooling primarily results in the loss of true signals rather than the introduction of spurious ones. These findings demonstrate that small pool sizes do not necessarily bias results, but instead reduce statistical power, leading to underestimation of the extent of biological change.

Consistent with this interpretation, effect size comparisons showed that while direction and magnitude of diet effects were largely preserved across pool sizes, estimates from n = 5 were substantially noisier and less reliable than those from larger pools. This increased variability was concentrated among metabolites with moderate and small effect sizes, whereas metabolites with large diet effects remained relatively stable even at small pool sizes. Together, these results indicate that small pools may be sufficient to detect the strongest signals, but are inadequate for capturing the full spectrum of biologically relevant variation.

### Pool size and replication jointly determine signal detection

Replicate downsampling further revealed that both pool size and biological replication interact to determine the robustness of metabolite detection. Across all datasets, reducing replicate number led to a progressive loss of significant metabolites, with smaller pool sizes accelerating this decline. Notably, the loss of signal under downsampling was driven primarily by a reduction in true positives rather than an increase in false positives, reinforcing the conclusion that sampling limitations primarily affect power. Importantly, pool size strongly modulated the rate of signal loss: larger pools retained diet-associated signals across a broader range of replicate reductions, whereas pool size = 5 exhibited rapid degradation of both statistical significance and effect size stability. These results demonstrate that pool size and replication are not interchangeable, but instead play complementary roles in determining the reliability of metabolomic inference.

The robustness of metabolite detection was also strongly dependent on effect size. Metabolites with large diet effects were consistently retained across downsampling conditions, whereas those with smaller effects were rapidly lost as replication and pool size decreased. While this pattern was consistent in direction across datasets, the rate at which signal was lost varied substantially, indicating that dataset-specific properties influence the sensitivity of metabolite detection.

### Signal detection is governed by biological effect size and variability

To identify the factors underlying these differences, we modeled metabolite detection frequency as a function of effect size, variability, pool size, and replication. Retention was strongly positively associated with effect size and negatively associated with variability in effect estimates, indicating that metabolites with stronger and more stable signals are more robust to reduced sampling. These results demonstrate that signal detection is governed by a balance between biological signal strength and measurement variability, with experimental design modulating this relationship. Differences in robustness across datasets are therefore explained not only by sampling design, but also by variation in the underlying signal-to-noise structure of the metabolome.

At the level of biological interpretation, functional module analysis revealed that not all metabolic pathways are equally robust to reduced sampling effort. Lipid-associated pathways and one-carbon metabolism showed relatively high retention across downsampling conditions, whereas redox metabolism and nucleotide turnover were more sensitive to reductions in pool size and replication. These differences suggest that some biological processes generate stronger or more coherent metabolomic signals, making them more resilient to experimental limitations. Importantly, this demonstrates that sampling decisions influence not only the number of detected metabolites, but also which biological processes are ultimately inferred to be affected.

### Implications for metabolomics experimental design

Together, these findings have important implications for experimental design in metabolomics. First, very small pool sizes (e.g., n = 5) should be avoided when the goal is to detect moderate or subtle biological effects, as they substantially reduce sensitivity and increase variability in effect size estimates. Second, increasing pool size from 5 to 50 individuals provides a large improvement in both profile stability and signal detection, whereas further increases from 50 to 100 yield diminishing returns. Third, biological replication remains critical across all conditions, and reductions in replicate number lead to consistent losses in detectable signal regardless of pool size. These results indicate that optimal experimental design requires balancing pool size and replication, rather than prioritizing one at the expense of the other.

Several limitations of this study should be considered. Our analyses were conducted in *Drosophila melanogaster* using whole-body metabolomics, and the extent to which these findings generalize to other organisms, tissues, or targeted metabolomics approaches remains to be determined. In addition, pool sizes were evaluated at discrete levels (5, 50, and 100), and intermediate values may provide further insight into the shape of these relationships. While we observed clear differences in signal robustness across datasets, our results indicate that these differences arise from variation in both effect size distributions and measurement variability among metabolites. This suggests that the detectability of metabolite signals is not solely determined by experimental design, but also by intrinsic properties of the measured metabolome, including the magnitude and stability of underlying biological effects.

## Conclusion

In conclusion, our results demonstrate that pooling is not a neutral technical decision, but a key determinant of both metabolomic measurement and biological inference. Pool size and replication jointly shape the detectability, stability, and interpretability of metabolomic signals, with their effects interacting in a nonlinear and dataset-dependent manner. As metabolomics continues to expand in evolutionary and systems biology, careful consideration of sampling design will be essential for ensuring robust and reproducible insights into the molecular basis of biological variation.

## Supporting information

Supplementary Figure 1. Per-metabolite pool-size sensitivity.

## Data availability

All code and supplemental materials used in this study are publicly available at GitHub (https://github.com/hubertdl/Metabolomics_sampling_project_D.melanogaster). Processed data and intermediate files necessary to reproduce all analyses and figures are included in the repository. Raw metabolomics data will be made publicly available in an appropriate repository prior to publication.

