## Supplementary figures and images for "Signal, noise, and sampling: How pool size and replication shape metabolomic inference"

### Supplementary Figure 1. Per-metabolite pool-size sensitivity.

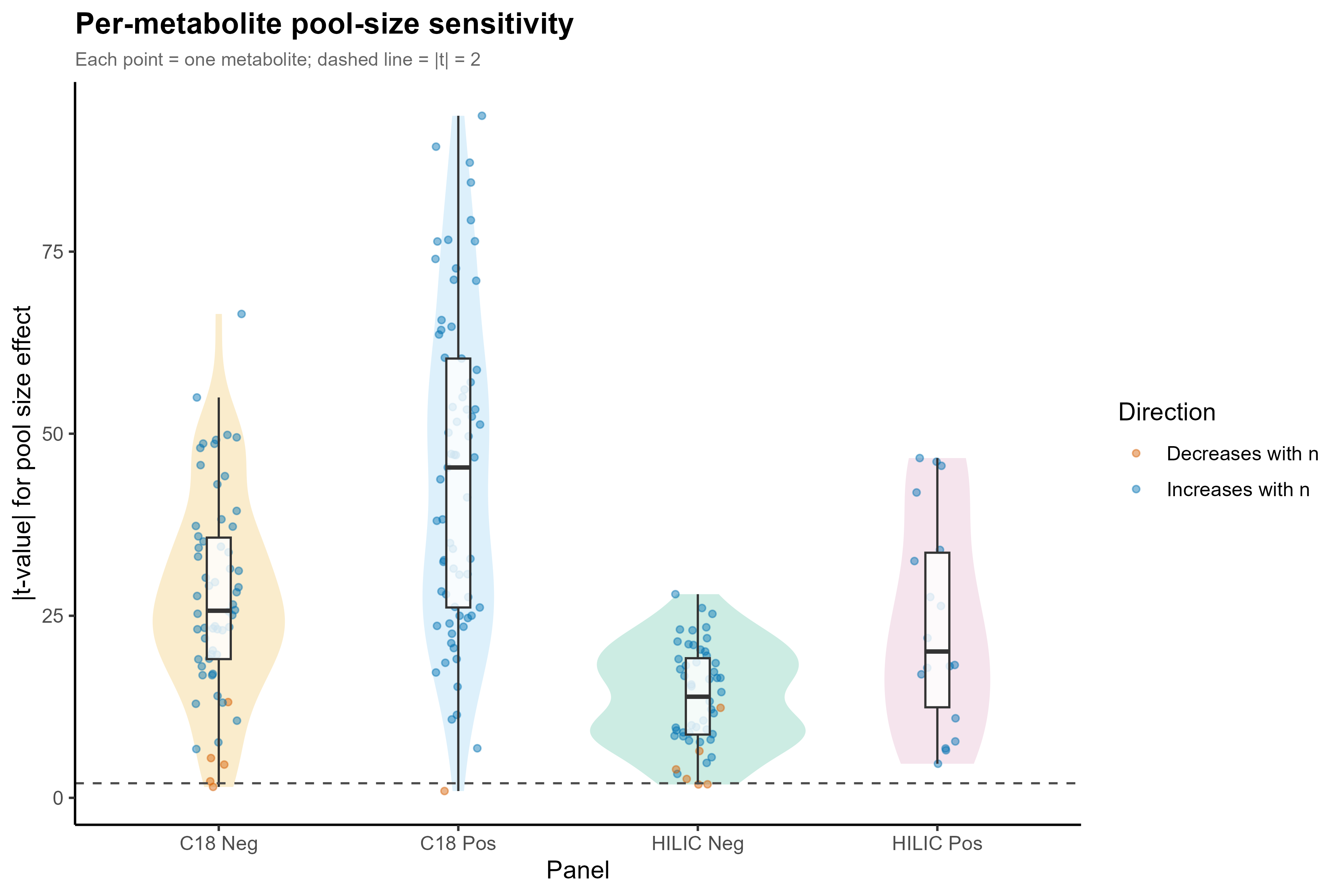
